# CoBeL-RL: A neuroscience-oriented simulation framework for complex behavior and learning

**DOI:** 10.1101/2022.12.27.521997

**Authors:** Nicolas Diekmann, Sandhiya Vijayabaskaran, Xiangshuai Zeng, David Kappel, Matheus Chaves Menezes, Sen Cheng

## Abstract

Reinforcement learning (RL) has become a popular paradigm for modeling animal behavior, analyzing neuronal representations, and studying their emergence during learning. This development has been fueled by advances in understanding the role of RL in both the brain and artificial intelligence. However, while in machine learning a set of tools and standardized benchmarks facilitate the development of new methods and their comparison to existing ones, in neuroscience, the software infrastructure is much more fragmented. Even if sharing theoretical principles, computational studies rarely share software frameworks, thereby impeding the integration or comparison of different results. Machine learning tools are also difficult to port to computational neuroscience since the experimental requirements are usually not well aligned. To address these challenges we introduce CoBeL-RL, a closed-loop simulator of complex behavior and learning based on RL and deep neural networks. It provides a neuroscience-oriented framework for efficiently setting up and running simulations. CoBeL-RL offers a set of virtual environments, e.g. T-maze and Morris water maze, which can be simulated at different levels of abstraction, e.g. a simple gridworld or a 3D environment with complex visual stimuli, and set up using intuitive GUI tools. A range of RL algorithms, e.g. Dyna-Q and deep Q-network algorithms, is provided and can be easily extended. CoBeL-RL provides tools for monitoring and analyzing behavior and unit activity, and allows for fine-grained control of the simulation via interfaces to relevant points in its closed-loop. In summary, CoBeL-RL fills an important gap in the software toolbox of computational neuroscience.

## 1 Introduction

The discovery of place cells in the hippocampus (O’Keefe and Dostrovsky, 1971) and grid cells in the medial enthorinal cortex (Hafting et al., 2005) have advanced our knowledge about spatial representations in the brain. These discoveries have also spurred the development of computational models to complement electrophysiological and behavioral experiments, and to better understand the learning of such representations and how they may drive behavior (Bermudez-Contreras et al., 2020). However, the computations performed during spatial navigation have received far less attention. In recent years, reinforcement learning (RL) (Sutton and Barto, 2018) has been of increasing interest in computational studies as it allows for modeling complex behavior in complex environments. Reinforcement learning describes the closed-loop interaction of an agent with its environment in order to maximize rewarding behavior or to minimize repulsive situations. A common way of solving the RL problem consists of inferring the value function, a mapping from state-action pairs to expected future reward, and selecting those actions which yield the highest values. The brain is thought to support this value-based type of RL (Schultz et al., 1997), and RL has been used to explain both human (Redish et al., 2007; Zhang et al., 2018) and animal behavior (Bathellier et al., 2013; Walther et al., 2021) (see Botvinick et al. (2020) for a recent review).

Concurrent with advances in understanding neural spatial representations, our theoretical understanding of reinforcement learning has also improved significantly in recent years. One major innovation was the combination of RL with deep neural networks (DNNs) (Mnih et al., 2015). These Deep RL models are now among the best-performing machine learning techniques, reaching higher performances than humans on complex tasks like playing Go (Mnih et al., 2015) or video games (Silver et al., 2016, 2017).

These advances in RL methods offer the opportunity to study the behavior, learning, and representations that emerge, when complex neural network models are trained on tasks similar to those used in biological experiments. The use of virtual reality in animal experiments (Pinto et al., 2018; Koay et al., 2020; Nieh et al., 2021) would even allow for the direct comparison of *in-vivo* and *in-silico* behavior. First attempts in this direction have recently shown that Deep RL models develop spatial representations similar to those found in enthorinal cortex (Cueva and Wei, 2018; Banino et al., 2018) and hippocampus (Vijayabaskaran and Cheng, 2022).

In many computational studies, models are custom built for specific experiments and analyses. While several software frameworks for simulations have been developed, they are often specialized for a certain type of task (Eppler et al., 2009; Leibo et al., 2018). Furthermore, the majority of frameworks are developed primarily within the context of machine learning and for eventual practical applications (Beattie et al., 2016; Chevalier-Boisvert et al., 2018; Liang et al., 2018), which makes their adaptation for neuroscience a daunting task. The high heterogeneity of models with respect to their technical design, implementation and requirements severely complicates their integration. A common RL framework would provide an opportunity to share and combine work across different fields and levels of abstraction.

Here, we introduce the ”*Closed-loop simulator of **Co**mplex **Be**havior and **L**earning based on **R**einforcement **L**earning and deep neural networks*”, or *CoBeL-RL* for short. The framework is based on the software that had been developed for studying the role of the hippocampus in spatial learning (Walther et al., 2021; Vijayabaskaran and Cheng, 2022) and further extends and unifies its functionality. The CoBeL-RL framework offers a range of reinforcement learning algorithms, e.g. Q-Learning or deep-Q network, and environments commonly used in behavioral studies, e.g. T-maze or Morris water maze. In contrast to other libraries, virtual environments across different levels of abstraction are supported, e.g. gridworld and 3D simulation of physical environments. Furthermore, our framework provides tools for the monitoring and analysis of generated behavioral and neuronal data, e.g. place cell analysis.

## 2 Methods

The CoBeL-RL framework provides the tools necessary for setting up the closed-loop interaction between an agent and an environment Sutton and Barto (2018). CoBeL-RL focuses on the simulation of trial-based experimental designs, that is, the learning of a task is structured into separate trials, similar to behavioral experiments. Single trials are (usually) defined by the completion of the task or reaching a timeout condition, i.e. time horizon. Each trial is further differentiated into agent-environment interactions referred to as steps. Each step yields an experience tuple (*s_t_, a_t_, r_t_, s*_*t*+1_), which the agent can learn directly from or store in a memory structure for later learning. *s_t_* is the agent’s state at time *t*, *a_t_* is the action selected by the agent, *r_t_* is the reward received for selecting action *a_t_*, and *s*_*t*+1_ is the agent’s new state.

CoBeL-RL is separated into modules that can be classified into three categories (Agent, Environment and Utility), which provides models of the RL agent, models of the environment, and utility functionalities, respectively (Fig.1). These three categories of modules will be explained in detail in the following sections. The majority of the framework is written in Python 3 (Rossum, 1995), while a few components are written in other programming languages as required by the software to which they interface. Tensorflow 2 (Abadi et al., 2015) serves as the main library for implementing Deep Neural Network models, and Keras-RL2 (Tensorflow 2 compatible version of Keras-RL (Plappert, 2016)) as the base for the framework’s deep Q-network (DQN) agent (Mnih et al., 2015). The interaction between RL agents and RL environments is facilitated through Open AI Gym (Brockman et al., 2016). PyQt5 is used for visualization.

**Figure 1.**
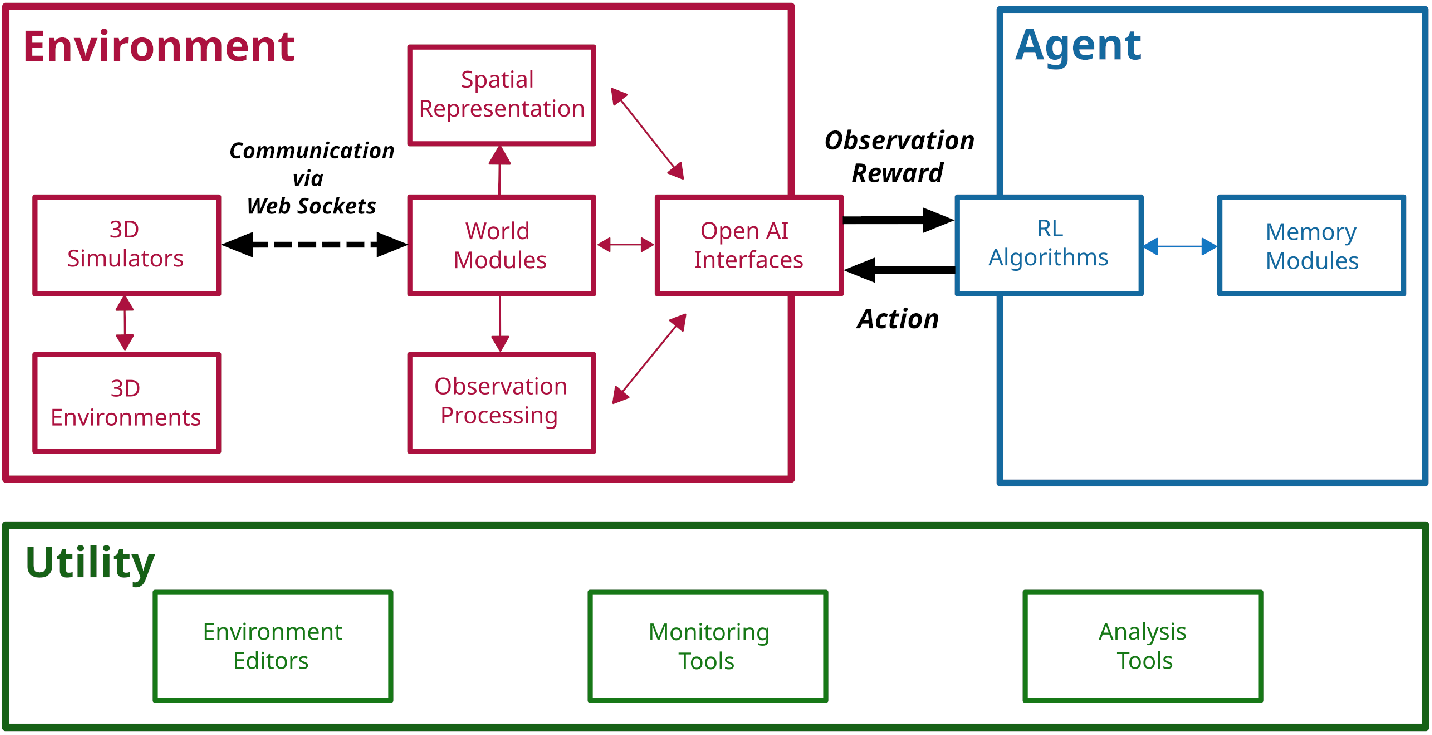
The CoBeL-RL framework. Illustration of CoBeL-RL’s modules and their roles. Modules can be grouped into three categories: Modules, which handle the simulation of the virtual environment and the flow of information (Environment), implement different RL algorithms (Agent), and implement supporting functionalities (Utility) such as monitoring and data analysis. The interactions between an agent and its environment Sutton and Barto (2018) is implemented with the help of the module ”OpenAI Interfaces” and provides observation and reward information to the agent, and send action to be executed to the environment.

### 2.1 Agent Modules

RL agents are implemented via the Agent modules that defines the behavior including learning, exploration strategies, and memory (see Fig. 1). All RL agents inherit from a common abstract RL agent class requiring them to implement functions for training and testing, as well as the computation of predictions for a given batch of observations. Furthermore, all RL agents implement callbacks which are executed at the start and end of a trial or step to allow for fine-grained control during simulations. Trial information, e.g. number of trial steps, reward, etc., is collected by RL agents and passed to the callbacks. All callback classes inherit from a common callback class, and custom callback functions can be defined by the user and passed as a dictionary to the RL agent.

RL agents in CoBeL-RL can use experience replay Lin (1992a,b); Mnih et al. (2015) as part of the learning algorithm. To do so, agents are linked with different memory modules used internally as buffers for experience replay. Experience tuples are stored in the memory modules providing the possibility of building up a history of experiences, which are used for training. Agents and memory modules can be combined freely to study the effects of different replay models.

Four agents are currently available and described briefly in the following.

#### 2.1.1 DQN Agents

The framework’s baseline DQN agent encapsulates Keras-RL2’s DQN implementation. It uses a small fully connected DNN by default and follows an epsilon-greedy policy. Since the original implementation is trained for a given number of steps instead of trials, the aforementioned callbacks are used to allow training for a given number of trials.

This agent’s callback class inherits from both the Keras callback class as well as the common callback class. Furthermore, versions of the DQN which implement Prioritized Experience Replay (Schaul et al., 2016) (PER-DQN) and learn an environmental model are also available.

#### 2.1.2 Dyna-Q Agents

The Dyna-Q model (Sutton and Barto, 2018) is implemented as a static tabular agent, that is, the agent’s Q-function is represented as an array of size |*S*| × |*A*| where |*S*| is the number of environmental states and |*A*| is the number of available actions. Due to its tabular nature, this agent, and those that derive from it, can only be used in conjunction with discrete static environments that represent states as abstract indices, such the gridworld interface. The agent’s environmental model is encapsulated as a separate memory module and similarly represents the environment in a tabular fashion. The memory module stores and retrieves experiences. For action selection either an epsilon-greedy (default) or softmax policy can be chosen. Furthermore, an optional action mask can be used to remove actions from the action selection that do not result in a state change. The agent’s Q-function is updated each step online and via experience replay (Lin, 1992b). Experience replay can be performed after each step, after each trial, or disabled. In addition, an implementation of the Prioritized Memory Access (Mattar and Daw, 2018) model is also provided.

#### 2.1.3 Dyna-DQN and DSR Agents

CoBeL-RL further offers hybrid agents which are derived from the Dyna-Q agent and use a DQN to represent their Q-function which we refer to as Dyna-DQN. The DQN part of the hybrid does not rely on Keras-RL2 and can be implemented separately in Tensorflow 2 or PyTorch (Paszke et al., 2019). An abstract network class serves as an interface between the agent and the separately implemented network. A set of observations which correspond to the different discrete environmental states can be passed to the agent. If no observations are defined, a one-hot encoding of the environmental states is generated which serve as observations. Additionally, a hybrid agent that learns a version of the Deep Successor Representation (DSR) (Kulkarni et al., 2016) is also provided which we refer to as Dyna-DSR. Similar to the DQN agents, the Dyna-DSR uses a small DNN by default to represent the Q-function. However, unlike the DSR, the Dyna-DSR does not learn a separate feature representation of its observations. Instead, reward and successor representation models are trained directly on the observations.

#### 2.1.4 Model-Free Episodic Control Agents

While all three Q-learning-based agents introduced above update the Q-function incrementally, Model-Free Episodic Control (MFEC) (Blundell et al., 2016) agents are designed for fast learning by repeating the best action they have performed in a specific state in the past. MFEC is therefore well suited for modeling one-shot learning (Wiltgen et al., 2006; Kosaki et al., 2014). The Q-function of MFEC agents is represented by an array of growing size where a state-action pair and its corresponding Q value is inserted if the pair is encountered by the agent for the first time. The Q value is updated in a one-shot manner, which simply keeps the best accumulative reward encountered so far. During inference, the agent searches the array, finds the most similar state to the current state and retrieves the Q values upon which action selection decisions are made. To further improve computational efficiency, all unique, high-dimensional states are first projected to a low dimensional space and then stored by using a KD-tree data structure (a K-dimensional tree is a binary tree where every node is a k-dimensional point) (Bentley, 1975), so that the search of closest neighbors to a given state (measured under Euclidean distance) becomes efficient.

### 2.2 Environment Modules

The implementation of the RL environments is split across different modules, and the agents interact with them through Open AI Gym interfaces. Roughly, CoBeL-RL provides two types of environments: simple environments that are directly implemented within an interface, e.g. a gridworld, and complex environments that are rendered by a game engine, such as Unity (Unity Technologies, 2005), Godot (Linietsky and Manzur, 2007), or Blender Game Engine (Blender Online Community, 2018). The latter involves additional modules for interaction with game engines (3D Simulators), processing of observations (observation modules) and navigation (spatial representation modules).

#### 2.2.1 Interface Module

All interfaces inherit from the Open AI Gym interface (Brockman et al., 2016) and implement step and reset functions. CoBeL-RL offers a wide range of different interfaces.

The gridworld interface represents gridworld environments in a tabular fashion. Environmental transitions are determined via a state-action-state transition matrix of size |*S*| × |*A*| × |*S*|. |*A*| = 4 to realize the movements on the grid (up, down, left, right). The reward function and terminal states are represented as tables of size |*S*|, where the reward function is real valued and the terminal states are binary encoded. The set of possible starting states is represented as a list of state indices.

A simple 2D environment is implemented by the 2d interface and allows interaction either by moving in the four cardinal directions or as a differential wheeled robot (i.e. the agent can move either of its wheels). In the former case the agent’s state is represented as 2d coordinates and in the latter case orientation is also included. The environment itself has no obstacles with the exception of walls that delineates the area in which the agent can navigate. The environment can contain multiple reward locations. Trials end when the agent reaches a reward, and optionally when it hits a wall.

The discrete interface implements tabular environments similar to the gridworld interface but is compatible with more complex environments. Like the gridworld interface, it requires the definition of a state-action-state transition matrix, a reward function, a table containing terminal states, and a list of starting states. Unlike the gridworld interface, it provides observations, which can be defined for each state, instead of state indices. If no observations are defined, a one-hot encoding is generated for each state.

The baseline interface provides a simple interface implementation for deep Q-Learning agents. The baseline interface defines the interaction of the baseline DQN agent with the spatial representation module, the observation module and the simulated 3D environment. The number of actions available to the agent is determined by the spatial representation module. The following sections detail the functionality of spatial representation, observation and 3D simulator modules.

#### 2.2.2 Spatial Representation Module

The spatial representation module allows the agent to navigate on a simplified spatial representation of the environment, rather than continuously through space. Currently, this module constructs a topological graph of the environment, with nodes and edges defining the connectivity. The topological graph can either be manually defined by the user when working with Blender by placing nodes and edges directly in the Blender Game Engine (Blender Online Community, 2018) (BGE), or can be automatically constructed using the grid graph module. CoBeL-RL provides implementations of simple rectangular and hexagonal graphs, and other graph types, such as a Delaunay triangulation, can be easily implemented if needed.

The spatial representation module can also be used to directly define the action space of the agent and specify how the actions are mapped to transitions on the graph. The default topological graphs implement two modes of transitions - without rotation, which only allows transitions to neighboring nodes, and with rotations, which allows both translational and rotational movements on the graph.

#### 2.2.3 Observation Module

The observation module provides functionality for the pre-processing of observations retrieved by the environment before it is sent to the RL agent.

The simplest observation module retrieves the agent’s position in x-y coordinates as well as its current heading direction, which can be used by the RL agent alongside or instead of visual observations. The observation module also pre-processes visual observations by resizing them to a user-defined size and normalizing the pixel values in the range [0, 1], such that it can be passed to the agent. Furthermore, observations can be corrupted with different types of noise, e.g. Gaussian noise, to better capture the imprecision of biological observations. Two or more observation modules can also be flexibly combined to simulate multisensory observations. For multisensory simulations, the individual observation modules are stored in a dictionary. The observations are then passed to the RL agent through the interface module. For deep RL agents, the keys of the dictionary can be used to indicate the input layer of the neural network to which those observations should be passed, allowing the use of complex network structures to process the observations.

#### 2.2.4 3D Simulators

The 3D Simulators module implements the communication between the CoBeL-RL framework and game engines that are used for simulation and rendering. The sending of commands and retrieval of data is handled via web sockets. CoBeL-RL supports three different game engines for simulation and rendering: Blender Game Engine (BGE) (Blender Online Community, 2018), Unity (Unity Technologies, 2005), and Godot (Linietsky and Manzur, 2007).

The baseline BGE Simulator module communicates via three separate web sockets. The control socket is used to send commands and retrieve control relevant data, e.g. object identifiers. Commands are encoded as strings which contain the command name as well as parameter values in a comma-separated format. Similarly, retrieved values arrive as strings in a comma-separated format. Observational data like visual observations and sensory data are retrieved via the video socket and data socket, respectively. During module initialization, a new Blender process is launched and a user-defined Blender scene is opened. Important commands provided by the module include the changing of an agent’s position and orientation, as well as the control of light sources. Observational data is automatically retrieved when the agent is manipulated, but can also be retrieved manually.

The Unity Simulator module is based on the *Unity ML-Agents Toolkit* (Juliani et al., 2020), which provides various interfaces for the communication between Unity and Python. The toolkit is mainly divided into two parts, one for the Unity side and another one for the Python side. The former is written in *C*#, which is the programming language for developing Unity games. Data transmission is handled by web sockets in the *Unity ML-Agents Toolkit*. Users do not need to set up the sockets themselves, but rather use the APIs provided by the Toolkit. Both string and float variables can be exchanged between the two sides.

The Godot Simulator module is built on the Godot engine and follows the same communication scheme as the BGE Simulator module. However, instead of Python, *GDScript* is used by Godot. Furthermore, instead of using comma-separated strings for communication most data is encoded in the JSON format. The Godot Simulator module supports the same commands and functionality as the BGE Simulator module and additionally allows for the switching of environments without the need for restarting the 3D simulator.

### 2.3 Utility Modules

CoBeL-RL’s utility modules consist of the analysis module and misc module. Simulation variables, e.g. behavior and learning progress, can be monitored using the analysis module’s various monitor classes. Gridworld tools and an environment editor can be found in the misc module.

#### 2.3.1 Analysis Module

The analysis module contains code for different types of analysis and monitoring tools.

CoBeL-RL provides monitors which store variables of interest during the course of training: the number of steps required to complete a trial (escape latency), the cumulative reward received in each trial, the responses emitted by the agent on each trial or step, and the agent’s position at each step (trajectory). Additionally, for Deep RL agents, the activity of layer units can be tracked with the response monitor. The different monitors can be added to any RL agent and are updated with trial information from callback calls.

The analysis tools enable the computation of spatial activation maps of network units at a desired resolution. The maps can be generated during the course of training or at the end of training by artificially moving the agent on a rectangular grid in the environment and recording the activity of the units of interest at the grid points. Based on the spatial activation maps, CoBeL-RL also provides a method to identify units that show special spatial firing properties, such as place-cell-like units and vector-like units (Vijayabaskaran and Cheng, 2022).

#### 2.3.2 Misc Module

The misc module provides tools that support the setting up of simulations and environments.

The gridworld tools provide functionality for the creation of gridworld environments. Gridworlds can be generated by either manually defining the relevant variables, e.g. size, reward function, starting states, etc., or by using templates for specific instances of gridworld environments, e.g. an open field with a single goal location. The gridworld tools generate the variables required by the gridworld interface, such as the state-action-state transition matrix, reward function, etc., and store them in a dictionary. A visual editor for gridworlds, which builds on the gridworld tools is provided in the gridworld GUI.

The Unity editor tools allow the user to quickly design the structure of a discrete, maze-like environment including external walls, obstacles, an agent and rewards, and import the generated file to the Unity editor to create a virtual environment. The tools are written in Java (version 17.0.1) and include a GUI as back-end functionalities, which generate the .yaml file required by the Unity editor. The user can choose from a large number of textures, and has the option of adding and deleting materials.

### 2.4 Additional Details About Offline Rendering Experiment

In Section 3.3 we show two examples of simulations using the BGE and Unity simulators. For the online/offline rendering experiment in Sec. 3.3, we use a T-maze environment with a Gridworld topology. The scene consisted of 3,061 vertices and 2,482 faces, including a visual representation of the agent that was partly visible in every frame and a single spherical light source. Maze walls had image textures and used diffuse (Lambert) and specular (CookTorr) shading. Backface culling was active for the complete scene. Simulations were run on a 16 core Intel(R) Xeon(R) W-1270P CPU with clock speed 3.80 GHz and 32GB RAM and a NVIDIA Quadro RTX(R) 5000 GPU. The agent was trained for 5 trials on the random pellet chasing task to encourage exploration of the entire maze, with a maximum trial duration of 400 time steps. For the offline rendering, observation images were pre-rendered and stored for all possible agent locations and later retrieved when needed. Simulation time for pre-rendering and storage was excluded from the mean frame time analysis.

## 3 Results

### 3.1 CoBeL-RL

The CoBeL-RL framework is a highly modular software platform to conduct spatial navigation and learning experiments with virtual agents. Figure 2 illustrates one of its realizations to simulate a T-maze experiment where an agent has to visit the right arm of the maze to receive a reward. The representation of this environment was split into two parts: the visual appearance of the environment and the topology. The former was built using the Unity simulator, which renders RGB images and sends them to the observation module. These image observations can be resized, or normalized to conform to parameter ranges of pixel values and are then sent to the OpenAI Gym Interface. The topology module generates a topological graph of the environment automatically and sends the spatial information of the artificial agent (blue square in Fig. 2) to the Gym interface as well. The reward function was defined inside the Gym interface to decide what kind of behavior is reinforced. For example, if the reward (green ball, Fig. 2) is placed in a fixed location in the maze in every trial, then a simple goal-directed navigation task is simulated.

**Figure 2.**
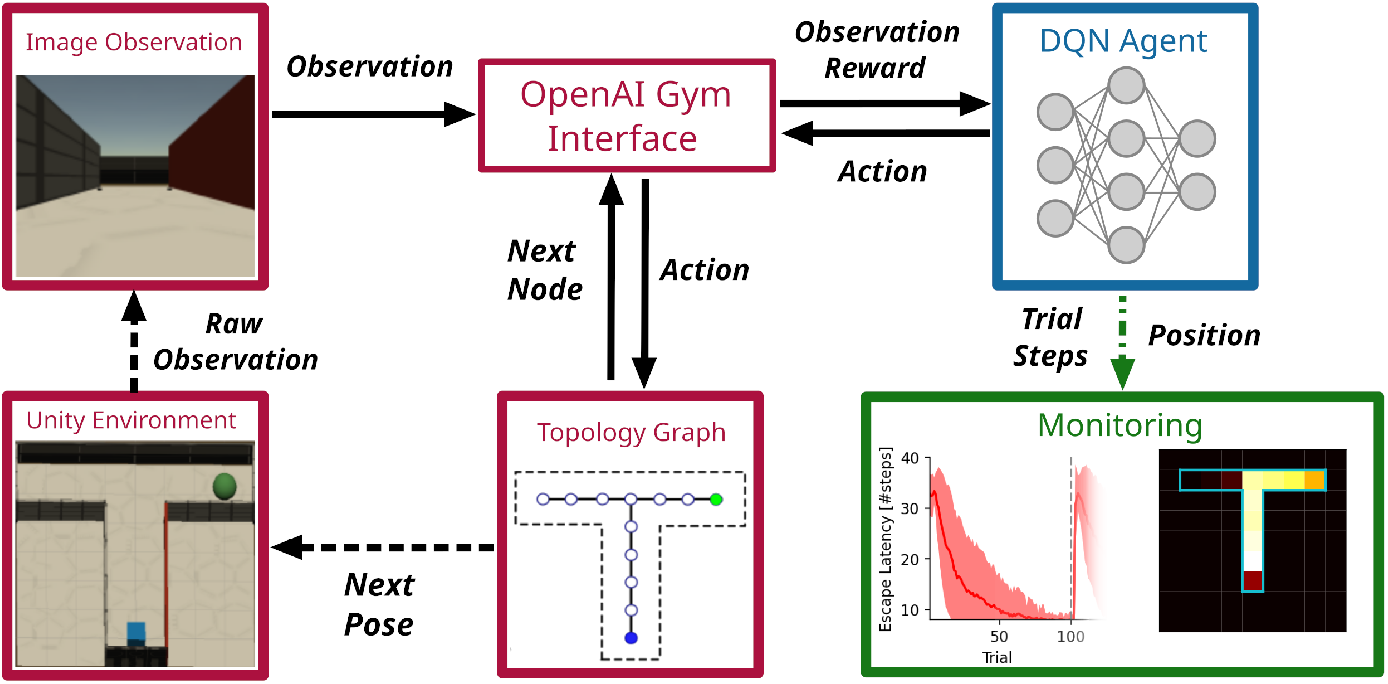
The CoBeL-RL framework in use. Example of simulation loop employing a DQN agent in a virtual 3D environment with visual stimuli. Simulation relevant variables are passed between CoBeL-RL’s modules directly (solid arrows) and via web sockets between CoBeL-RL and the 3D simulator (dashed arrows). The monitor modules rely on callback to record variables of interest (dashed green arrow).

As part of the Gym Interface, a DQN agent implemented using Keras/Tensorflow receives the processed image observation, the instant reward and then takes an action, deciding whether the artificial agent should move forward, turn left, etc. The DQN agents can utilize memory replay to speed up learning and to study the role of short-term memory and episode replay in spatial navigation tasks (Zeng et al., 2021). It is possible to turn the memory on or off, limit its size, or change the statistics of how memories are replayed to model the effect of manipulating episodic memory (Diekmann and Cheng, 2022). The action signal is further passed from the Gym interface to the topology graph, where the position and orientation of the artificial agent is computed and sent to the Unity simulator to update the virtual environment, which closes the feedback loop between agent and environment. At the same time, the performance, i.e. escape latency, cumulative response curve, etc., of the DQN agent is monitored live and displayed to the user.

### 3.2 Fast and Easy Creation of Environments

We evaluated CoBeL-RL on a set of environments that model biological experiments with rodents. CoBeL-RL provides multiple options to set up a virtual environment. The requirements for the set-up differ based on the type of virtual environment: Gridworld environments require the definition of a transition graph, while 3D environments further require the modeling and texturing of the environment’s geometry. In addition to these fully manual methods, CoBeL-RL provides simple visual editors for the quick creation of maze environments at different abstraction levels, i.e. 2D gridworld environments and 3D environments in the Unity Game Engine. As an example we demonstrate how both of them can be used to set up a simple T-maze environment in the following sections.

#### 3.2.1 Gridworld Editor

The Gridworld editor provides an intuitive graphical interface for the creation of gridworld environments. After first selecting the height and width of a new gridworld, the editor presents a grid of states which can be interacted with using the mouse (Fig. 3A). States can be selected via a mouse left click and their properties edited, e.g. the reward associated with a state. Transitions can be edited by double clicking on edges between adjacent states, thereby toggling the transition probability between the states in both directions between zero and one. For more fine grained control, state-action transitions can be manually defined upon selecting the advanced settings option in the properties of a currently selected state. Gridworlds created with the editor are stored using Python’s *pickle* module and directly used with CoBeL-RL’s gym gridworld and gym discrete interfaces. The gridworld itself is represented as a state-action-state transition matrix, reward function, starting set (i.e. the states at which an agent may start), and terminal set (i.e. the states at which a trial terminates). Additionally, metrics like the number of states, state coordinates, and a list of invalid transitions are also stored.

**Figure 3.**
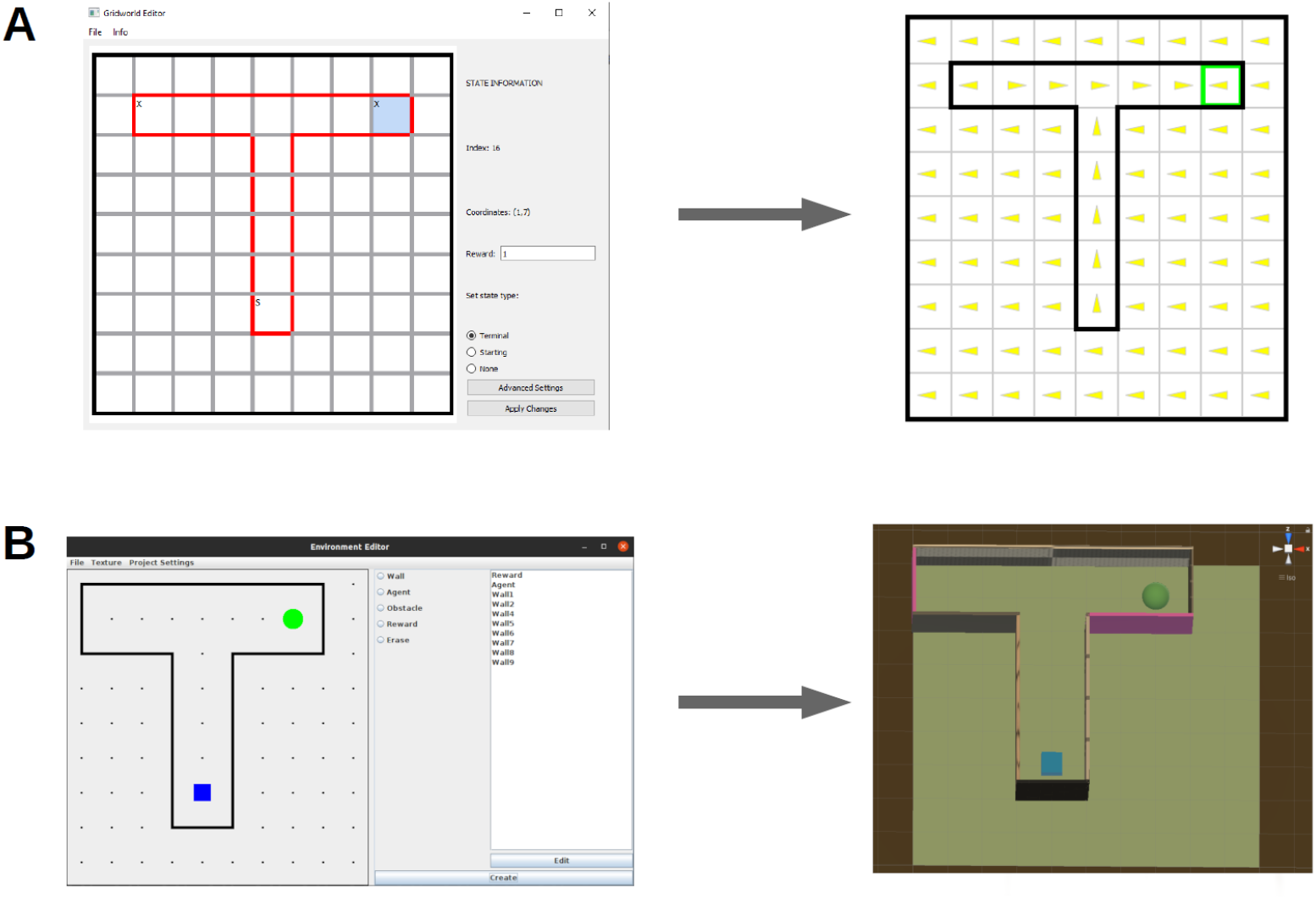
World editors. **A:** The gridworld editor (left) can be used to quickly and intuitively create new environments for simulation (right). **B:** The editor for 3D Unity environments (left) generates 3D Unity scenes (right).

#### 3.2.2 Unity Editor

The Unity Editor can be used to easily build 3D maze environments, which can then be employed as the training and testing environments for the RL agents in CoBeL-RL. An interactive canvas covered with grids in the GUI (Fig. 3B) allows the the user to simply draw the positions and dimensions of the maze walls as well as the locations of obstacles, an agent and reward(s). The grid size corresponds to one unit length in Unity and the dimension of the entire 2D grids are adjustable. Therefore, the user is able to design mazes of any size and shape. The editor has included a library of different texture materials. After the structure of the maze has been defined, the user can choose to either manually select a material from the library for each object in the maze, or let the editor randomly assign a texture for that same object. The files for all materials are placed in a folder, and the user has the option to add or delete any material from within the GUI. Finally, the Unity scene files can be directly imported into the Unity Editor and start communicating with the CobeL-RL framework via the frontend interface for Unity.

### 3.3 Efficient Distributed Simulation Using CoBeL-RL

CoBeL-RL can be used in a closed-loop simulation to simulate the interaction of the agent and its environment online. Using the renderer online in a closed-loop setup has the advantage of being very flexible and reflecting changes in the environment directly. However, often the environment tested for simple RL agents only consists of a small set of possible observations. For instance, in the Gridworld example in Fig. 2, the agent can be located in one of only 12 different positions. Even if multiple configurations, like heading direction or different visual cues are allowed for every location, the total number of possible image observations will still be small compared to the number of learning iterations. In such situations, it may be beneficial to resort to an offline rendering strategy, i.e. to render and store the image observations once at the beginning of the simulation and then retrieve the observations from memory instead of rendering them anew in every time step. This mode is crucial when simulations are run on a remote compute cluster that does not have the required software packages installed to run the renderer locally. CoBeL-RL provides the functionality to switch between online and offline rendering with little changes to the simulation code. In this section we provide a comparison between these two modes in terms of the simulation runtime.

For a numerical comparison, we used a simple T-maze environment that was rendered using Blender and a foraging task to encourage exploration using the MFEC agent Blundell et al. (2016) (see Methods for details). For online rendering, Blender was queried to retrieve image observations at every simulation time step. For the offline rendering case, this was only done once for all possible image observations at the beginning of the simulation. Images were then stored in the main memory for later retrieval during the simulation. Frame time was measured by invoking the system clock at the beginning of every simulation time step. As expected, average frame times over the entire simulation were much higher for online than for offline rendering (Fig. 4).

**Figure 4.**
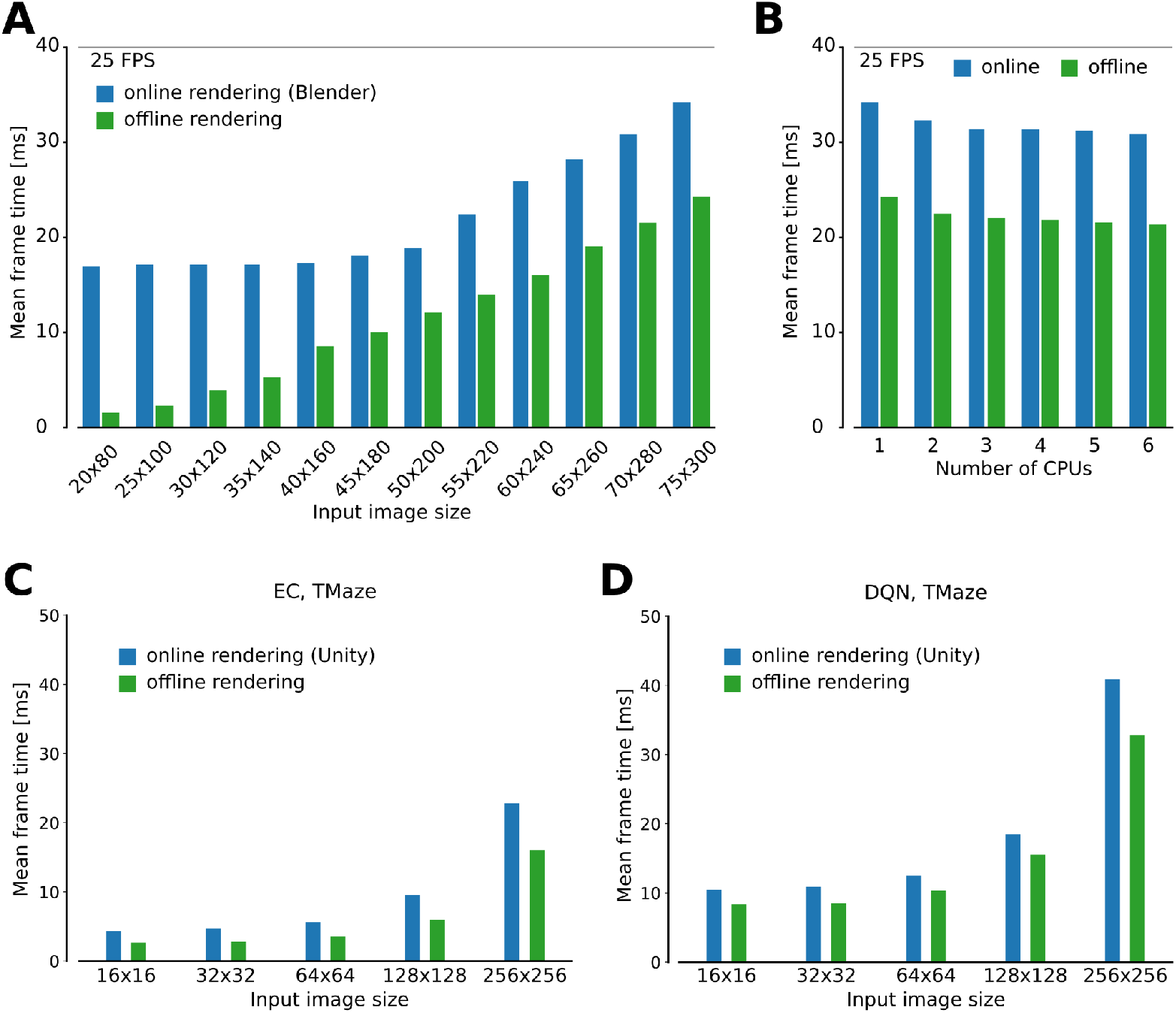
Analysis of simulation speed. **A:** Comparison of simulation time of a single step using the online and offline rendering method with Blender for observations of different size. **B:** Scaling of run time with number of CPUs. **C,D:** As in (A) but for the Unity rendering instead of Blender, using the MFEC (A) and DQN (B) agent.

The speedup gained by offline rendering is likely to depend on a number of factors, of which we explore the image size, the number of CPU cores, game engine and RL algorithm. First, the simulation time is expected to strongly depend on the size of the image observations, we tested different sizes between 20×80 and 75×300 pixels and confirmed that the relative overhead for rendering varied greatly with the image size (Fig. 4A). Specifically, we found that for small input image sizes the overhead of rendering with Blender was substantial. For instance, for the smallest tested image sizes of 20×80, online rendering frame time was 16.9 ms, more than eleven times the frame time for offline rendering, which was 1.5 ms. For larger input images this difference was significantly less pronounced, e.g., for the 75×300 pixel images, mean frame time was 34.1 ms and 24.2 ms for online and offline rendering, respectively, a factor of only ca. 1.4. Second, somewhat surprisingly increasing the number of CPU cores beyond two did not have a strong impact on the simulation speed for both online and the offline rendering (Fig. 4B). We hypothesize that this is due to the low level of parallelism that can be achieved in the small models that we studied here. Third, scaling of frame time with image size was qualitatively similar for the Unity renderer (Fig. 4C,D). Finally, we compared two different RL agents, MFEC and DQN, where the latter is more computationally expensive than the former because of the need for GPU computing. The relative overhead for rendering increases for both agents when the image size expands. On the other hand, although the total frame time for the DQN is larger than that for MFEC, the relative overhead for rendering is comparable between the two agents for all image sizes (Fig. 4C,D). This result shows that the contribution of the agent and the renderer to the total simulation time, are modular and do not much interfere with one another.

In summary, we find that offline rendering is especially beneficial if small image observations are used since the overhead for rendering dominates the simulation in this situation. Frame time speedups are achieved with offline rendering for all frame sizes, but this method is only suitable if the total number of possible image observations is small.

### 3.4 Analysis of Behavior With CoBeL-RL

To analyze the behavior of the simulated agents, relevant parameters have to be recorded and evaluated. To do so, CoBeL-RL provides various monitors with which relevant variables can be recorded, visualized and evaluated during training (Fig. 5). Monitors are available for tracking trial reward, escape latency, and cumulative responses. For the last monitor, the specific response coding can be defined by the user. The default coding is one when reward was collected in a trial, and zero if not. These three behavioral performance monitors are compatible with all agents. Below we demonstrate the use of CoBeL-RL’s performance monitors in two example experimental paradigms.

**Figure 5.**
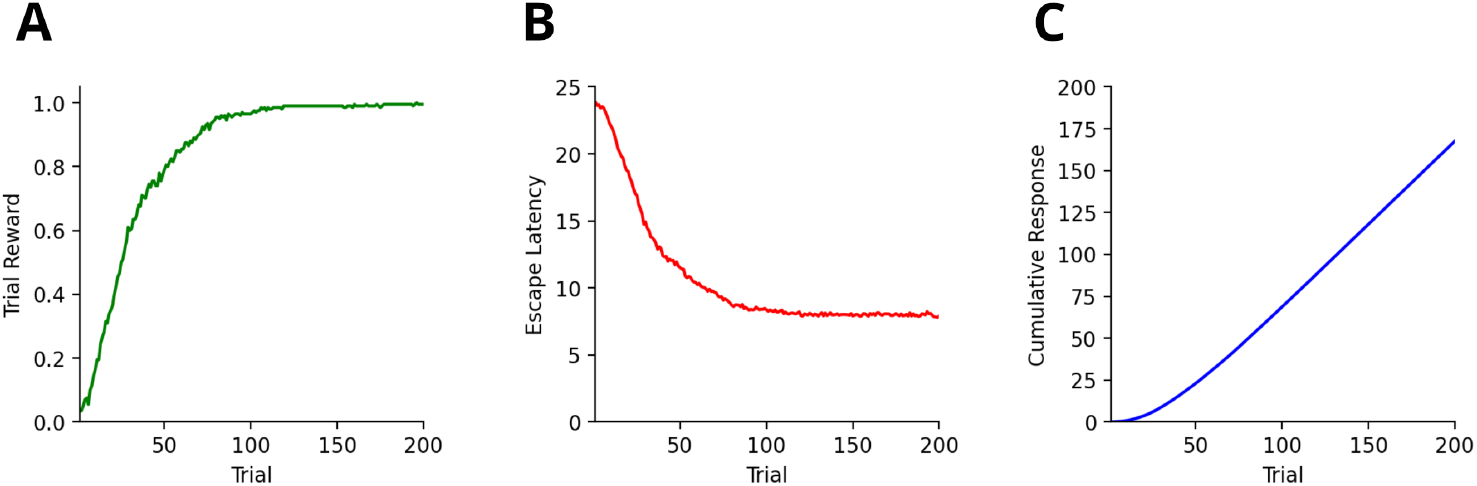
Commonly needed monitors in CoBeL-RL. **A**: The reward monitor tracks the cumulative reward collected during each trial. **B**: The escape latency (in number of trial steps) can be tracked with the escape latency monitor. **C**: The response monitor can track the cumulative responses emitted by the agent. Per default responses are coded with one when a reward was collected and zero otherwise. The coding of responses can be customized by using the simulation loop’s callback functionality.

The analysis of responses emitted by an agent is important to understand the reinforcement of behaviors in classical and instrumental conditioning. We therefore simulated a simple extinction learning paradigm: The agent was placed in a T-maze environment and rewarded for choosing the right arm during the first 100 trials. Then reward was moved to the left arm for the remaining 100 trials. Responses are visualized by the cumulative response curves, where each trial with the choice of the right arm was encoded as one and the choice of the left (or neither) arm as zero, for a DQN agent in a complex 3D environment with visual stimuli (Fig. 6A) and for a tabular Dyna-Q agent in a simple gridworld (Fig. 6B).

**Figure 6.**
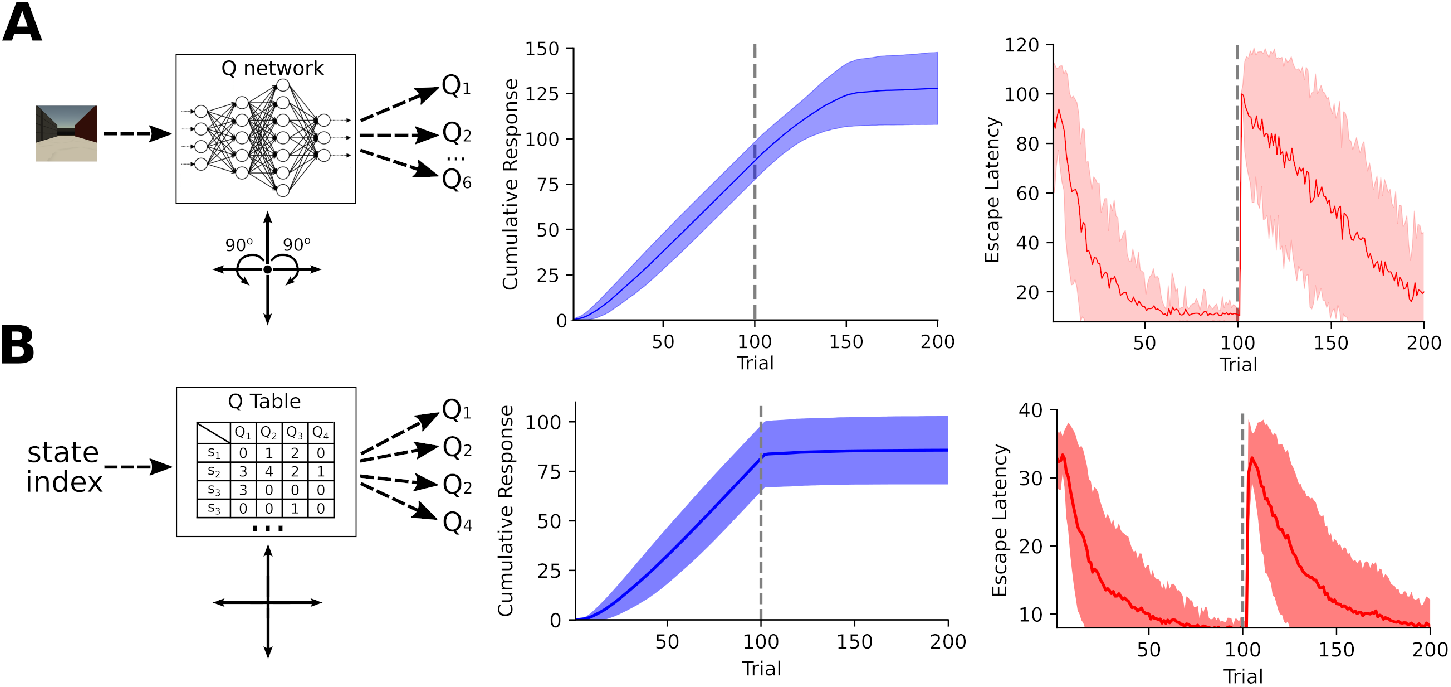
Analysis of behavior in an extinction learning paradigm. **A**: A DQN agent was trained on a simple extinction paradigm within a complex 3D Unity environment (left). The agent has six actions at its disposal: movement in either of the four cardinal directions and rotating either left or right by 90°. The cumulative response (center) reveals that the agent reliable picks the rewarded right arm for the first 100 trials. After the right arm was no longer rewarded (trials 101-200), the previously learned behavior initially persists and then gradually extinguishes. The escape latency curves (right) show that agent quickly learned to reach the right arm. After the reward switched the agent slowly adapted to the new reward location. **B**: Same as **A** but recorded from a Dyna-Q agent (left) in a gridworld environment. Unlike the DQN agent the Dyna-Q agent can only move in the four cardinal directions. Similar to the DQN agent the Dyna-Q agent reliably picks the rewarded right arm for the first 100 trials. However, due to the Q-function’s tabular representation the previously learned behavior extinguishes rapidly.

Another example is latent learning in the Blodgett maze Blodgett (1929), a composition of 6-Unit Alley T-Maze (Fig. 7A), which we simulated using the Dyna-DSR agent in CoBeL-RL (see Methods for details). Agents were tested in two settings: latent learning, in which the environment was devoid of reward for the first 100 trials (exploration phase; –100 to –1 trials) and reward was introduced for the remaining trials (Fig. 7B, purple line.), and direct learning, in which there was no exploration phase and reward was present from the first trial in the maze (Fig: 7B, orange line). Latent learning agents learned to reach the goal state faster than direct learning agents after the reward was introduced. These results qualitatively reproduce experimental findings Blodgett (1929); Reynolds (1945); Tolman (1948).

**Figure 7.**
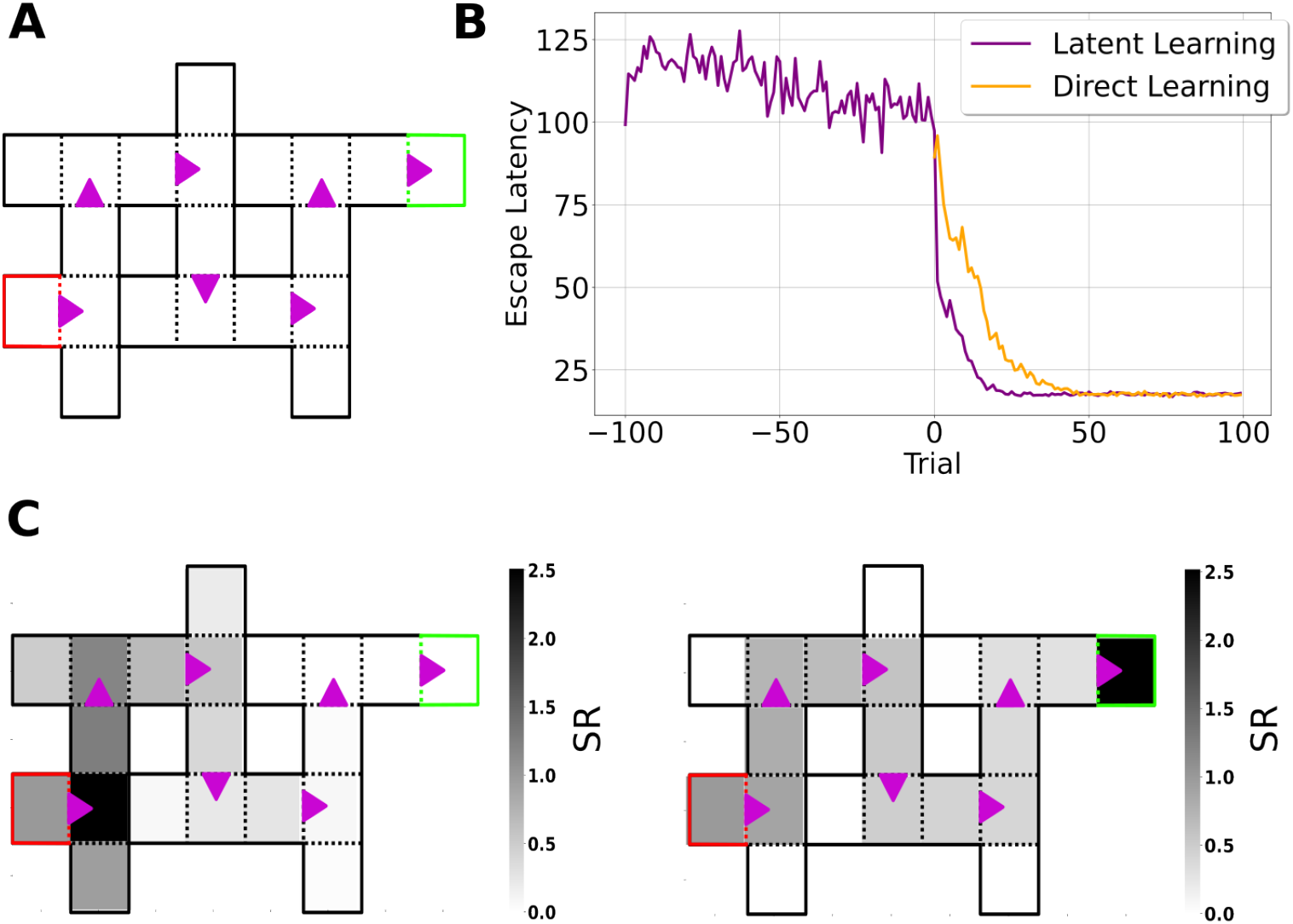
Simulation of latent learning using CoBeL-RL. **A:** Gridworld of the Blodgett maze. The red and green squares represent the start and goal states, respectively. Purple arrows indicate transitions that are only allowed in the indicated direction. **B:** Escape latency of agents trained in latent learning (purple) and direct learning (orange) settings. For better comparison, the curve for the direct learning setting was shifted to the point of reward introduction of the latent learning setting. Curves represent means over 30 simulations per agent. **C:** Learned Deep SR of the start state at the end of the training phase (left) and at the end of the simulation (right). At the end of the training phase, the learned SR mainly represents states close to the start state. In contrast, by the end of the simulation, the learned SR represents the path to the goal state. The color bar represents the learned cumulative discounted (state) occupancy.

### 3.5 Analysis of Neural Representations With CoBeL-RL

An important goal of CoBeL-RL is to allow the analysis of not only behavioral measures of the reinforcement learning agents, but also the computations and representations that emerge in the deep neural network to support behavior. To this end, CoBeL-RL provides response monitors which can track network representations of Deep RL agents. In the latent learning simulations above, we analyzed the learned Deep SR of the *start* state to examine what the agent had learned. At the end of the training phase (Fig. 7C, left), the learned SR mainly represents states close to the start state, simply reflecting the environment’s topology. In contrast, by the end of the simulation (Fig. 7C, right) the learned SR represents the path to the goal state.

CoBeL-RL also includes analysis tools to analyze spatial representations, including the ability to compute spatial activity maps, classify them, and identify units in the network that have place fields. To demonstrate this functionality, we built a simple Blender environment (Fig. 8A) and trained the agent to solve a goal finding task on a hexagonal topology graph (Fig. 8B). In each trial, the agent had to navigate to the unmarked goal node (indicated in green) from a random starting position. The agent had twelve available actions – six translations, and six rotations. The translations allowed the agent to move to any of the six neighboring nodes on the topological graph, and the rotations turned the agent in place to face a neighboring node. We generated spatial activity maps by placing the agent in all locations of a 25 × 25 grid that was embedded in the environment, which can be set flexibly depending on the desired resolution in CoBeL-RL, and recorded the activation of nodes in the DQN network. Place field maps of all units preceding the output action selection layer of the Deep Q-Network were recorded every 800 trials during the training process. Recordings were obtained for all six heading direction of the agent and further processed to identify units that exhibit place-cell-like firing (Fig. 8C) – see 2 for a detailed description. As would be expected, if place-field-like representations supported spatial navigation, the number of neurons that exhibit place fields increased during training (Fig. 8D). In general, the spatial activation maps can be constructed either at set points during the training process to understand how they evolve with learning, as in this example, or after the training is completed.

**Figure 8.**
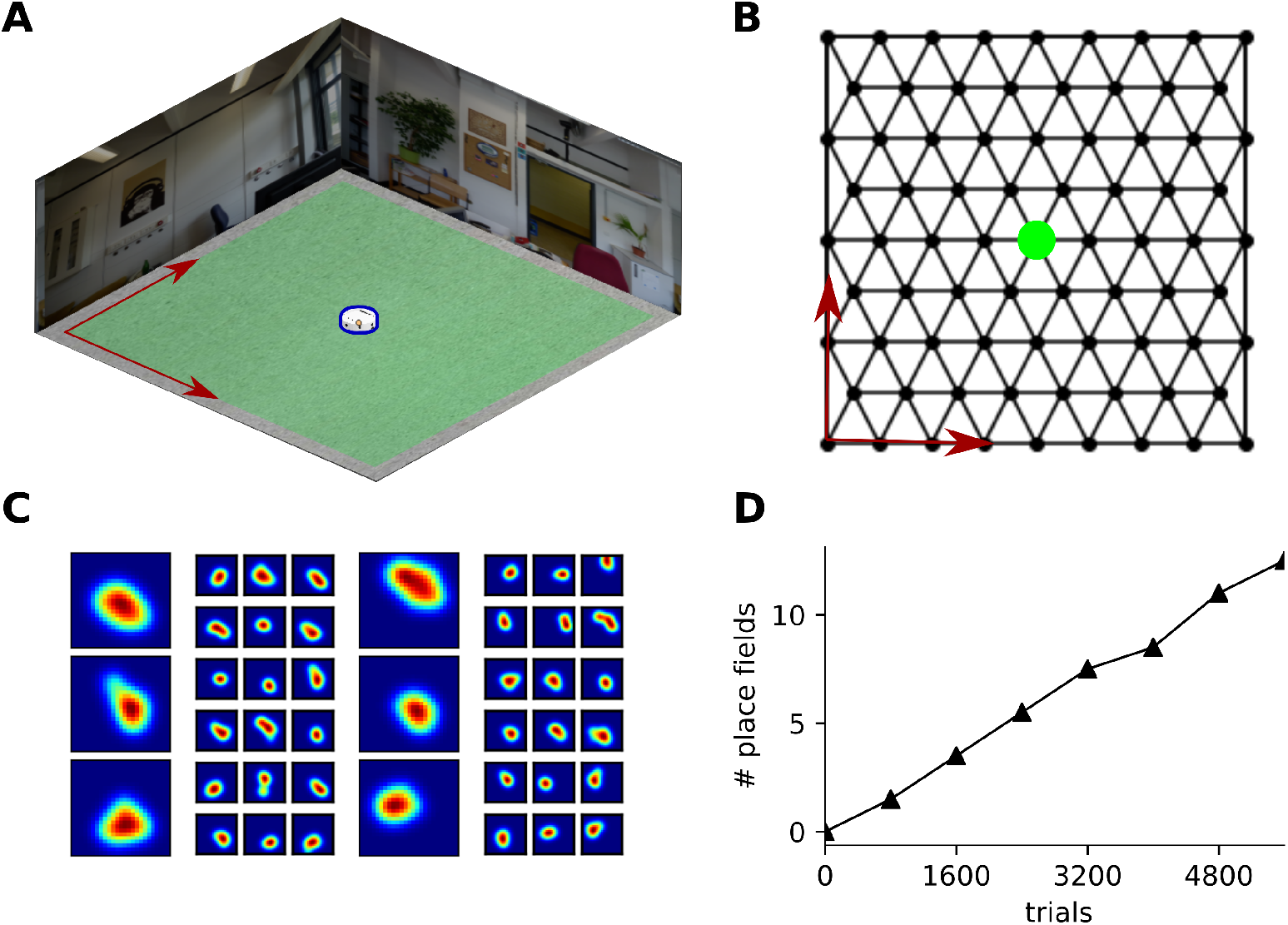
Analysis of spatial represenations using CoBeL-RL. **A:** Blender environment used for simulating a navigation task. The agent is marked in blue. **B:** Topology graph on which the agent navigates. The location of the unmarked goal is indicated in green. **C:** Example place cells emerging in the simulation. The large maps show the place field activity averaged over all head directions and the adjacent six smaller maps show the corresponding activity at each head direction, spaced 60 degrees apart. **D:** Evolution in number of place fields with learning trials.

## 4 Discussion

In this paper, we introduced CoBeL-RL, a RL framework oriented towards computational neuroscience, which provides a large range of environments, established RL models and analysis tools, and can be used to simulate a variety of behavioral tasks. Already, a set of computational studies focusing on explaining animal behavior (Walther et al., 2021; Zeng et al., 2021) as well as neural activity (Diekmann and Cheng, 2022; Vijayabaskaran and Cheng, 2022) have employed predecessor versions of CoBeL-RL. The framework has been expanded and refined since these earlier studies. Here, we provided additional details about the simulation framework and how it can be extended and used in future work.

### 4.1 Related Work

A number of RL simulation frameworks have been introduced previously. Most of them target machine learning or industrial applications. For example, DeepMind Lab provides a software interface to a first-person 3D game platform to develop general artificial intelligence and machine learning systems (Beattie et al., 2016). RLlib is a Python-based framework that provides a large number of virtual environments and toolkits for building and performing RL simulations (Liang et al., 2018). The main targets of RLib are industrial applications in domains such as robotics, logistics, finance, etc. One unique advantage of RLlib is that it offers architectures for large-scale distributed RL simulations to massively speed up training. The framework also supports Tensorflow and Pytorch. RLib has been used by industry leaders, but does not target scientific studies in neuroscience. The Minimalistic Gridworld Environment (MiniGrid) provides an efficient gridworld environment setup with a large variety of different types of gridworlds, targeting machine learning (Chevalier-Boisvert et al., 2018). Gridworld environments are based on OpenAI Gym and could therefore be easily integrated with CoBeL-RL. MiniGrid differs from our Gridworld interface in that it provides isometric top view image observations instead of abstract states. MAgent comprises a library for the efficient training of multi-agent systems Zheng et al. (2018); Terry et al. (2020) and includes implementations of common Deep RL algorithms like DQN.

On the other hand, there are efforts that are targeted towards application in neuroscience. For instance, SPORE provides an interface between the NEST simulator, is optimized for spiking neuron models Eppler et al. (2009), and the GAZEBO^1^ robotics simulator (Kaiser et al., 2019). SPORE targets robotics tasks, and the framework is not easy to use for analysis of network behavior or emerging connectivity in the network. The framework also cannot be easily combined with deep Q-learning. Psychlab aims at comparing the behaviors of artificial agents and human subjects by recreating the experimental setups used commonly in psychological experiments (Leibo et al., 2018). These setups include visual search, multiple object tracking and continuous recognition. The setups are recreated inside the DeepMind Lab virtual environment Beattie et al. (2016). RatLab is a software framework for studying spatial representations in rats using simulated agents (Schönfeld and Wiskott, 2013). The place code model is based on the hierarchical Slow Feature Analysis (SFA) (Schönfeld and Wiskott, 2015). RatLab supports a number of different spatial navigation tasks and can be flexibly extended. It is not well suited to be integrated with reinforcement learning methods.

### 4.2 Limitations and Future Developments

CoBeL-RL provides an efficient software framework for simulating closed-loop trial-based learning, which is well matched for modeling behavioral experiments. Nevertheless, experiments that do not have a strict trial-based structure or well-defined time horizon could be modeled in CoBeL-RL as well by controlling the training loop via CoBeL-RL’s callback functionality, although such simulations would be less efficient. In addition, CoBeL-RL’s monitoring tools would require adaptation to account for the potentially undefined time horizon and lack of trial structure.

The experimental paradigms discussed here only include one agent. However, the effect of another agent’s presence on behavior (Boesch and Tomasello, 1998; Kruützen et al., 2005) and neural activity (Rizzolatti and Craighero, 2004; Mukamel et al., 2010) has also been of great interest. Currently, CoBeL-RL does not support simulations that involve multiple agents and at first glance it is not clear how they could fit in the trial-based training loop. However, such multi-agent simulations could be facilitated by extending CoBeL-RL with an abstract supervisor agent which can encapsulate an arbitrary number of agents. The training loop of this abstract supervisor agent could then control the training of the agents it supervises. The behavior of the agents could be easily monitored by creating separate instances of the monitoring classes for each agent.

Currently, CoBeL-RL largely relies on the Tensorflow-2-based implementations of Deep RL agents provided by Keras-RL2. This strong reliance on Tensorflow 2 could be detrimental to the appeal and longevity of CoBeL-RL: Multiple programming libraries for the implementation of DNNs exist (Al-Rfou et al., 2016; Paszke et al., 2019) and change over time. Agents would have to be re-implemented for different libraries and updated whenever a library’s behavior changes. We address these problems with the use of an abstract network class which serves as an interface between Deep RL agents and specific network implementations. Currently, network classes for Tensorflow 2 and PyTorch are supported.

As a framework, CoBeL-RL is continuously developed and extended. While current efforts focus on simulations in virtual environments, CoBeL-RL can be connected to physical robots like the Khepera-IV (Tharin et al., 2019). Agents can be trained efficiently in simulation and then take control of the physical robot. Pretraining an agent in simulations before letting it control a real robot has been shown to work in other settings (James and Johns, 2016; Tzeng et al., 2017).

### 4.3 Conclusion

In conclusion, CoBeL-RL is an RL framework oriented towards computational neuroscience that fills an gap in the landscape of simulation software, which currently focuses mostly on machine learning, a specific task paradigm, or certain type of model. Importantly, CoBeL-RL provides a set of tools which simplify the process of setting up simulations through its environment editors. This is the case especially in the context of 3D simulations since otherwise their creation would require the user to acquire a wide range of additional skills, e.g. 3D modeling, game engine programming, etc. CoBeL-RL hence greatly reduces the overhead of setting up closed-loop simulations which are required to understand the computational issues that animals face in behavioral tasks and the solutions that they generate.

## Author Contributions

ND, SV, and SC contributed to conception and design of the framework. ND, SV, XZ, DK, and MM performed and analyzed simulations. ND, SV, XZ, DK, and MM contributed to the code. All authors contributed to the first draft of the manuscript, and edited and approved the submitted version.

## Acknowledgments

We thank Dr.-Ing. Thomas Walther for the conceptualization and development of CoBeL-RL’s initial version.

## Data Availability Statement

The CoBeL-RL framework is available at https://github.com/sencheng/CoBeL-RL. The simulation code used to produce the results is available at https://github.com/sencheng/CoBeL-RL-Paper-Simulations.

## Footnotes

1 http://gazebosim.org/

